# High-quality reference genome of the African hermit spider, *Nephilingis cruentata,* and sex chromosome evolution in spiders

**DOI:** 10.64898/2026.08.18.745515

**Authors:** Hans Recknagel, Elena Bužan, Luka Močivnik, Paul Vincent Debes, Cene Fišer, Carolina Ortiz-Movliav, Simona Kralj-Fišer

**Affiliations:** University of Ljubljana, Biotechnical Faculty, Department of Biology, Ljubljana, Slovenia; University of Trier, Department of Biogeography, Trier, Germany; University of Primorska Faculty of Mathematics, Natural Sciences and Information Technologies, Koper, Slovenia; Faculty of Environmental Protection, Velenje, Slovenia; Department of Aquaculture and Fish Biology, Hólar University, Sauðárkrókur, Iceland; Research Centre of the Slovenian Academy of Sciences and Arts, Institute of Biology, Ljubljana, Slovenia

**Keywords:** Nephilingis cruentata, chromosome-level assembly, sex chromosomes, Hi-C, Araneae, synteny

## Abstract

**Background:** Chromosome-level genome assemblies are increasingly enabling tests of chromosome evolution, conserved synteny, and sex chromosome conservation across diverse animal lineages, including spiders.

**Results:** Here, we present a chromosome-level genome assembly for the African hermit spider, *Nephilingis cruentata*, a species with extreme female-biased sexual size dimorphism and a cytogenetically inferred X₁X₂ sex chromosome system. The final Hi-C-assisted assembly spans 1.72 Gbp, with 99.5% of bases assigned to 13 pseudochromosomes, a scaffold N50 of 131.6 Mbp, and a BUSCO completeness score of 98.8%. We annotated 20,021 protein-coding genes, and repetitive elements accounted for 42.7% of the genome. Sex-specific whole-genome resequencing identified Chr02 and Chr07 as candidate X chromosomes based on reduced male coverage, consistent with the expected X₁X₂ system. Using comparative whole-genome alignments across existing chromosome-scale spider assemblies, we also show that sex-linked chromosomes retain broad homologous identity across sampled spider lineages but exhibit lower synteny conservation and greater chromosome-length divergence than autosomes.

**Conclusions:** These results suggest that spider sex chromosomes are conserved in homologous identity but more labile in structure, providing a comparative framework for studying sex chromosome conservation and divergence across Araneae.

## Introduction

Spiders (Araneae) are an ancient and diverse radiation of terrestrial predators comprising three major lineages: the segmented Mesothelae, the Araneomorphae, and the largely fossorial Mygalomorphae. They have colonized nearly every terrestrial habitat on Earth and show extensive variation in silk and venom biology, web architecture, dispersal strategy, reproductive biology, and life history [1–3]. This ecological and phenotypic diversity makes spiders a useful system for investigating how variations in genomes contributes to phenotypic evolution.

Understanding the genomic basis of this diversity requires chromosome-level genome assemblies that place genes, repetitive elements, duplicated regions, and structural rearrangements within their physical chromosomal context. Such assemblies enable analyses of conserved synteny, gene-family evolution, chromosome fusions and fissions, repeat-associated genome expansion, genetic mapping, and other large-scale genomic patterns that fragmented assemblies often cannot resolve [4–7]. This is particularly relevant for spiders, whose genomes are often large, repeat-rich, and heterozygous, making assembly and annotation challenging [6–8]. Nevertheless, the increasing availability of chromosome-level spider genomes has enabled comparative analyses of genome size, repeat content, chromosome number, and synteny conservation across Araneae [5–7,9–12]. However, these resources have only begun to be used to investigate how genomic architecture relates to phenotypic evolution and chromosome evolution across spiders.

Sex chromosome evolution is of particular interest in spiders because they exhibit an exceptional diversity of chromosome systems and sex-related traits. Cytogenetic studies have identified a wide range of sex chromosome configurations, including X₀, X₁X₂, X₁X₂Y, neo-sex chromosome systems, cryptic homomorphic chromosomes, and more complex multiple-X systems [6,13,14]. Among these, the X₁X₂ system (males: X₁X₂0, females: X₁X₁X₂X₂) is the most widespread sex chromosome configuration, particularly among araneomorph lineages and has been proposed as an ancestral condition for spiders [15,16]. The origin of the X₁X₂ system remains unresolved, with proposed mechanisms including fission of an ancestral single X chromosome [16,17] or nondisjunction of an X chromosome followed by differentiation of the newly formed sex chromosome [18]. Both mechanisms predict the formation of two non-homologous X chromosomes that subsequently follow distinct evolutionary trajectories. Variation in sex chromosome number and configuration has further been attributed to chromosome fission, fusion, nondisjunction, polyploidization, and sex chromosome–autosome rearrangements [7,13,14].

Sex chromosomes differ from autosomes in inheritance, effective population size, recombination environment, hemizygosity, and exposure to sex-specific selection, factors that can shape their evolutionary trajectories and structural differentiation [19–21]. These properties may also contribute to differences in the genomic basis of sex-specific development, sexual dimorphism, and reproductive traits. Despite extensive cytogenetic knowledge, the molecular basis of sex determination in spiders remains unresolved, and expanding genomic resources are needed to determine how sex chromosome configurations relate to their underlying genomic architecture [22]. Previous chromosome-level spider genomes have identified sex chromosomes in individual species and revealed conserved X-linked synteny among selected taxa, particularly in *Argiope bruennichi* and *Uloborus diversus* [6,23]. However, whether spider sex chromosomes retain homologous chromosomal identity across broader phylogenetic sampling, and whether chromosome-scale features of sex chromosomes differ from those of autosomes, remains incompletely resolved.

Repetitive DNA represents another important component of chromosome-scale genome organization because it contributes to genome size variation and chromosome structure. Transposable elements and other repeats can influence chromosome evolution through sequence expansion, heterochromatin formation, recombination-mediated rearrangements, and differences in chromosome size. In particular, repeat accumulation can contribute to sex chromosome differentiation in systems with suppressed or altered recombination [19–21]. In spiders, comparative analyses have linked genome-size variation to repeat content, particularly within Araneoidea [7]. However, whether variation in sex chromosome size reflects differential repeat accumulation compared with autosomes remains poorly understood in spiders.

The African hermit spider, *Nephilingis cruentata* (Fabricius, 1775), is a subtropical orb-weaving spider (Nephilidae, Araneoidea) distributed across sub-Saharan Africa and parts of South America [24]. The species shows extreme female-biased sexual size dimorphism, with females being approximately 75 times heavier and six times longer than males [2,25], underpinned by sex-specific developmental trajectories [26,27]. These pronounced sex differences in morphology and development, together with reproductive traits such as pedipalp breakage, mating plugs, and sexual cannibalism, make *N. cruentata* a biologically informative system for investigating how genomic architecture relates to sex-specific development and reproductive biology. Cytogenetic work on *N. cruentata* inferred an X₁X₂ sex chromosome system, with males carrying 2n = 24 chromosomes and females 2n = 26, but the sex chromosomes could not be distinguished morphologically from the autosomes [28]. *N. cruentata* therefore provides an ideal system for investigating the genomic features modulating sex-specific developmental and reproductive traits.

Previous work in the African hermit spider has shown that body size and growth are shaped by sex-specific genetic, maternal, developmental, and environmental effects. Male size is strongly influenced by developmental conditions and maternal effects [25,29], whereas female adult body size is more strongly associated with direct genetic effects [25]. These findings indicate that males and females may differ not only in phenotype but also in the genetic architecture underlying trait variation and responses to selection. Identifying the sex chromosomes is therefore necessary for distinguishing autosomal from X-linked genomic contexts, where inheritance, hemizygosity, recombination environment, and exposure to selection differ between males and females. A chromosome-level reference genome provides this framework and establishes the basis for testing whether genomic regions associated with sex-specific development, body size, and reproductive traits are autosomal, X-linked, or located in structurally divergent chromosomal regions.

Here we present the chromosome-level genome assembly of the African hermit spider, resolving 13 pseudochromosomes and identifying Chr02 and Chr07 as candidate X chromosomes based on depth-of-coverage analysis. Using chromosome-scale spider genomes spanning major lineages of Araneae, we test whether sex chromosomes retain homologous chromosomal identity across lineages and whether they differ from autosomes in synteny conservation and chromosome-length evolution. We further evaluate whether variation in genome size and total sex chromosome length is associated with genome-wide repeat content. Together, these analyses place the *N. cruentata* genome in a broader comparative framework for evaluating sex chromosome conservation, structural divergence, and chromosome-scale genome evolution in spiders.

## Methods

### Biological Material and Sampling

Specimens of *N. cruentata* were sourced from a long-term laboratory population maintained at the Institute of Biology, ZRC SAZU, Slovenia. The founding individuals originated from wild populations, collected in 2015 in iSimangaliso Wetland Park and Ndumo Reserve, South Africa. The specimens were laboratory-bred individuals from this long-term captive population. Due to inbreeding over multiple generations, such individuals may exhibit reduced genome-wide heterozygosity, which can facilitate genome assembly.

For genome sequencing, high-molecular-weight DNA was extracted from six adult females. The tissues used included the opisthosoma and prosoma, and DNA extracts from two females with the highest DNA quality were ultimately used for a genome survey via short reads and long-read sequencing (**Table S1**). Fresh specimens used for DNA extraction were flash-frozen in liquid nitrogen prior to extraction, processed according to the DNA extraction protocol, and stored at −80°C until further use.

For Hi-C library preparation, we used fresh tissue from four adult females, with chelicerae removed to reduce potential contamination from venom gland material (**Table S1**). Hi-C samples were flash-frozen in liquid nitrogen and processed following the Phase Genomics Proximo Hi-C Animal protocol, using DpnII as the restriction endonuclease supplied with the kit. Hi-C libraries were stored at −80°C until sequencing.

For transcriptome sequencing, we sampled spiders at different stages, i.e. fertilised eggs, hatchlings, juveniles, subadults and adults. Whole-body samples were used for hatchlings and juveniles before sex could be determined, whereas later developmental stages were sampled separately by sex and body region (details provided in **Table S2**). These samples were preserved in RNAlater, stored at −80°C, processed according to the RNA extraction protocol, and stored at −80°C until further use.

### DNA Extraction

Three kits were tested for DNA extraction: the DNeasy Blood & Tissue Kit (Qiagen), the MagAttract HMW DNA kit (Qiagen), and the Monarch HMW DNA Extraction Kit for Tissue (NEB). Out of these kits, the largest fragment sizes and concentrations were retrieved with the HMW DNA Extraction Kit for Cells & Blood (NEB), which was then used as a basis for DNA extraction for individual genome sequencing. DNA quality and quantity were assessed using Qubit dsDNA Broad-Range Assay Kit on a Qubit 3.0 Fluorometer (Thermo Fisher Scientific) and DNA integrity was assessed on a 0.8% agarose gel and a Tapestation 2200 with Genomic DNA ScreenTapes.

### Genome survey

Prior to long-read sequencing, a genome survey was conducted to obtain preliminary estimates of genome size, repeat content, heterozygosity, and GC-content. This was performed using ∼110 Gbases of short-read data generated from an MGI-2000 sequencer from a single female individual (**Table S1**). The genomic library was constructed using the MGIEasy Universal DNA Library Prep Kit V1.0 (CAT#1000005250, MGI) following the standard protocol. After sequencing, fastp [30] was used to filter raw reads (parameters: -n 0 -f 5 -F 5 -t 5 -T 5 -q 20). Cleaned reads were processed using KMC [31] (parameters: -k21 -ci1 -cs1000000) to generate the 21-mer frequency distribution. Next, genome size and heterozygosity were estimated with findGSE [32] and GenomeScope [33], and these estimates were used to inform the required long-read sequencing depth necessary for the genome assembly.

### Draft assembly with short and long reads

Following the short-read sequencing for the genome survey, DNA of a single female individual was used to construct a SMRTbell library for Pacbio Revio sequencing (**Table S1**). Quality control was performed in SMRT Link v12.0 removing failed reads (RQ<0.99). The initial genome assembly was generated with hifiasm v0.19 [34] using default parameters. Following the initial assembly, redundancy was removed based on pairwise alignment and collinearity among assembled sequences. Long, highly similar regions were identified using minimap2 v2.26 [43], considering alignments ≥10 kb with an edlib similarity ≥0.8. Contained redundant sequences were removed, while end-to-end overlapping sequences were merged to obtain a non-redundant preliminary assembly. A first evaluation of the assembly was performed with a BUSCO (Benchmarking Universal Single-Copy Orthologs) [35] search, relying on a set of conserved single-copy genes from the arachnida_odb10 dataset in the OrthoDB database (**Table S3**). Next, to determine the consensus quality value (QV) and K-mer completeness of the genome assembly, the clean read data was used to construct a 21-Kmer library using Merfin [36] and evaluated in Merqury v1.3 [37]. After this initial evaluation, the short reads generated for the genome survey were aligned to the preliminary assembly using bwa v0.7.17 [38] in mem mode with default parameters. Then samtools v1.18 [39], sambamba v1.0.0 [40], and freebayes v1.3.6 [41] (with default parameters) were used to read the single-base variant information and obtain a vcf file containing the variant results used to calculate the homozygous single-base variant rate. The single-base error rate of the genome was calculated by analysing homozygous and heterozygous mutation sites in the sample, with homozygous sites considered as genomic error sites. Next, cuteSV v2.0.3 [42] (parameters: --max_cluster_bias_INS 100 --diff_ratio_merging_INS 0.3 --max_cluster_bias_DEL 100 --diff_ratio_merging_DEL 0.3) was used to count the number of deletions (DEL), insertions (INS), inversions (INV), duplications (DUP), and breakend (BND) variants. Alignment of HiFi reads to the preliminary assembly was performed with minimap2, sambamba and samtools to identify coverage, heterozygosity, and GC content. Structural variation was inferred from pbsv v2.9.0 [44]. For the evaluation of contamination in the assembly, repetitive sequences were masked using RepeatMasker v1.331 [45] and aligned to the nucleotide sequencing (NS) database on NCBI using blastn. Prior to alignment, sequences above 1Mb were split into 50Kb bins and all sequences were then classified according to the NS database to estimate the degree of sequence contamination and to exclude contaminant sequences. Finally, after correcting the preliminary assembled genome sequences and conducting the de-contamination analysis, a BUSCO search was performed and the contig N50 statistics and genome size were recorded.

### Hi-C Library preparation, sequencing and chromosome-scale assembly

Hi-C libraries were prepared in-house from fresh tissue of an adult female using the Phase Genomics Proximo Hi-C Animal kit, with DpnII as the restriction enzyme. The prepared libraries were quality-controlled and sequenced by Novogene on a NovaSeq X Plus sequencing machine using 150-bp paired-end sequencing, generating 244.0 Gbp of raw sequence data.

Raw sequencing data was filtered in fastp v0.23.4 [30] (v0.23.4) to remove adapter sequences and low-quality reads. Next, seqkit v2.5.1 [46] was used to extract reads1 and reads2, and bowtie2 v2.3.2 [47] (alignment mode: --end-to-end; parameters: --very-sensitive -L 30) to perform a single-end alignment against the draft assembly. Unmapped reads were screened for junction sites (restriction site re-ligation), trimmed, and realigned. Then, PE reads from both alignment rounds were merged and the proportion of uniquely mapped PE reads was calculated. Next, HiC-Pro v3.1.0 [48] (default parameters) was used to perform statistical analysis of valid PE reads.

The quality-controlled reads were then aligned to the draft assembly using bwa-mem v0.7.17 [49] (parameters: mem −5SP), and bam-filter v2.0.0 (parameters: 1 --nm 3) was applied to filter for mapping quality and distance. Next, clustering, reassignment, ordering and orientation, and building pseudomolecules was run in HapHiC v1.0.5 [50] (command: Haphic pipeline Genome bam 13). The HapHiC pipeline corrects misassemblies, removes short and allelic contigs, clusters contigs using the MCL algorithm. Following initial clustering, contig order and orientation were manually curated based on Hi-C contact map signals. The curated contigs were then subjected to integrated 3D-DNA and ALLHiC optimization to determine contig order and orientation and generate chromosome-level pseudomolecules. Following initial clustering, the Hi-C contact maps were manually curated based on Hi-C heatmap signals to adjust the assembly before generating the chromosome-level pseudomolecules. The resulting Hi-C draft genome assembly was evaluated with N50 statistics and BUSCO analyses using Compleasm v0.2.5 [51].

### Transcriptome sequencing, assembly and annotation

RNA from 64 samples was extracted using the Qiagen RNeasy Micro Kit, including on-column DNase digestion to remove genomic DNA (**Table S2**). RNA integrity was assessed using an Agilent 2200 TapeStation. Libraries were prepared using the NEBNext Ultra II Directional RNA Library Prep Kit and sequenced on an Illumina NovaSeq 6000 system. Samples covered different developmental stages and both sexes to support genome annotation.

After sequencing, raw RNA-seq reads were filtered with fastp to remove adapter sequences, reads with an N content exceeding 10%, and reads in which more than 50% of bases had a quality score below 20. The filtered sequences were aligned to the draft assembly using STAR 2.7.3a [52]. Next, based on the mapped results, transcript assembly was performed using StringTie v1.3.4d [53]. The resulting transcript evidence was used in PASA v2.3.3 [54] to support genome-wide gene prediction and to generate a training set for ab initio gene prediction. Homology-based gene prediction was performed using GeMoMa v1.6.1 [55], using protein and annotation evidence from related spider species, including *Argiope bruennichi, Caerostris darwini, Nephila pilipes, Parasteatoda tepidariorum, Trichonephila clavata, Trichonephila clavipes,* and *Trichonephila inaurata madagascariensis*. For ab initio prediction, 3,000 high-confidence genes derived from transcript-supported gene models were selected to train a species-specific AUGUSTUS v3.3.1 [56] model. The trained model was then applied to perform *de-novo* gene structure prediction across the genome. Gene predictions from transcript-based, homology-based, and ab initio approaches were integrated using EvidenceModeler (EVM) v1.1.1 [54]. Evidence weights were set to prioritize transcript-supported predictions, followed by homology-based predictions and ab initio predictions (default weighting criteria). The integrated annotation produced the initial protein-coding gene set of the genome. Subsequently, genes containing transposable elements or encoding errors were filtered out via TransposonPSI [57], yielding the final gene set.

Non-coding RNAs (ncRNAs) were annotated using multiple approaches. Infernal v1.1.2 [58] was used together with the Rfam [59] database to identify structured ncRNAs. tRNAs were predicted with tRNAscan-SE v2.0 [60], and rRNA genes and subunits were predicted with RNAmmer v1.2 [61]. These results were integrated to generate the final ncRNA annotation for the genome.

Predicted genes were functionally annotated using multiple databases. BLASTp v2.7.1 searches were performed with an e-value threshold of 1e-5 and a maximum of one target sequence per query against the NR, KEGG, KOG, and SwissProt databases. KEGG was used for biological pathway annotation, KOG for eukaryotic orthologous group classification, and SwissProt for curated protein function annotation. Protein domains and gene ontology (GO) terms were inferred using InterProScan [62] with default parameters. Functional annotations from all databases were combined to produce the final annotation set.

### Repeat annotation

Microsatellite identification from the genome assembly was performed with GMATA v2.2 [63], and tandem repeats (TRs) were annotated with TRF v4.07b using default parameters. Prior to transposable element (TE) annotation, TR regions identified by TRF were soft-masked to reduce conflicts between tandem repeats and transposable elements. TEs were annotated using a custom repeat library generated from the TR-soft-masked genome. The custom library was built through a multi-step approach. First, MITE-Hunter [11] was used to identify miniature inverted-repeat transposable elements (MITEs). Next, the identified tandem repeats and MITEs were hard-masked with ambiguous bases (N) to facilitate de novo repeat identification using RepeatModeler v1.0.11 [45]. Repeats classified as “unknown” were further categorized using TEclass [64]. Finally, the constructed repeat libraries and the Repbase [65] database were integrated into a comprehensive reference library. This combined library was then utilized in RepeatMasker v1.331 to perform the genome-wide repeat annotation.

### Identification of candidate X chromosomes using sex-specific coverage

DNA from five male and five female individuals was extracted and sent for whole-genome sequencing to Novogene. Sequencing was carried out by Novogene on an Illumina NovaSeq X Plus platform using a paired-end 150 bp (PE150) strategy. Libraries were prepared by random fragmentation of genomic DNA, adapter ligation, size selection, and PCR amplification, with a target data output of approximately 17 Gb (10×) of raw data per sample (**Table S4**). Raw sequencing reads were assessed for quality using FastQC and processed using TrimGalore v0.6.10 (Phred score ≥5, minimum read length 75 bp, and removal of terminal ambiguous bases) to remove adapters and low-quality bases. Filtered reads were aligned to the chromosome scale reference genome using bwa-mem v0.7.19 [38]. Alignments were sorted and indexed, and the resulting BAM files were used as input for mosdepth v0.3.11 [66] to generate per-chromosome coverage summaries from genome-wide read depth, allowing the identification of candidate X chromosome chromosomes.

To account for differences in sequencing depth among individuals, coverage values were normalized within each sample before any comparisons across chromosomes or sexes were made. For each individual, we scaled each chromosome’s coverage by that individual’s overall genomic coverage level (the median across chromosomes). This ensures that differences in sequencing yield between individuals are comparable and that normalised coverage values reflect relative coverage within each individual genome, rather than absolute read depth. After this step, autosomes are expected to cluster around a common baseline close to one, whereas X chromosomes are expected to show reduced normalized coverage in males relative to autosomes and to females.

These normalized values were then used to (i) describe the distribution of chromosome-level coverage within males, allowing identification of the chromosomes with consistently reduced male coverage, and (ii) compare normalized chromosome-specific coverage between males and females by summarizing coverage separately for each sex and directly contrasting these summaries per each chromosome. Chromosomes showing consistent male-specific reductions in relative coverage were considered candidate X chromosomes.

### Comparative analyses of spider sex chromosome homology and chromosome evolution

Chromosome-scale spider genome assemblies were retrieved from public databases (N = 17; **Table S5**). Prior to whole-genome alignment, scaffolds not corresponding to assembled chromosomes or pseudochromosomes were excluded. To standardize repeat masking across assemblies, a species-specific repeat library was generated for each genome. RepeatModeler databases were first built using the BuildDatabase command in RepeatModeler v2.0.7 [67]. RepeatModeler was then run with the -LTRStruct option, except for *Ectatosticta davidi* and *Pardosa pseudoannulata*, for which this option was omitted because of technical issues. RepeatMasker v4.2.2 [68] was run with the -xsmall and -nolow options using the corresponding species-specific repeat libraries. BuildDatabase, RepeatModeler, and RepeatMasker were run from a containerized environment. RepeatMasker outputs were summarised and checked using a custom bash script.

Whole-genome alignments were generated with Progressive Cactus v3.1.3 [69] using the -logInfo option in a containerized environment. Synteny analyses were then performed from the resulting HAL alignments using halSynteny v2.2 [70]. Analyses were run pairwise, with each species iteratively treated as the reference and all other species treated as query species, one pairwise comparison at a time. halSynteny was run from the same containerized environment as Progressive Cactus.

To quantify chromosome-level synteny conservation, cactus alignment output was parsed to extract chromosome-level mapping summaries for all pairwise species comparisons. Each species assembly was iteratively treated as the reference genome against all other species, and for every reference chromosome the target chromosome with the highest alignment proportion was recorded. The resulting chromosome correspondence tables contained the strongest syntenic match for each reference chromosome and its corresponding chromosome-level synteny mapping proportion. These values were used to construct a weighted chromosome synteny network, where edge weights represented chromosome-level synteny mapping proportions between species.

To infer chromosome homology across all species, a weighted chromosome homology network was reconstructed from strong synteny links (mapping proportion ≥ 0.5), and homologous chromosome groups were identified using weighted Louvain community detection. These homology groups were then used to classify chromosomes as either autosomal or sex-linked. Sex chromosome conservation was assessed using two complementary classification schemes. First, a strict annotation approach considered only chromosomes explicitly identified as X chromosomes in species with experimentally characterized sex chromosomes. Second, a homology-based approach used these annotated X chromosomes as references to identify putative sex chromosomes in species lacking direct sex chromosome annotations. Specifically, chromosomes that clustered within the same chromosome homology groups as annotated X chromosomes were classified as putatively sex-linked.

Because homologous chromosomes are expected to retain chromosomal identity despite changes in sex chromosome system configuration, the composition of inferred sex-linked homologous groups was further examined across species. Specifically, we tested whether X chromosomes from different spider lineages consistently clustered within a single chromosome homology group, which would support retention of an ancestral X chromosome identity. Conversely, assignment of X chromosomes to multiple independent homology groups would be consistent with repeated recruitment of different chromosomes into sex chromosome systems.

A species-level maximum-likelihood phylogeny was reconstructed from a concatenated amino-acid alignment of 2,908 single-copy genes from the 17 spider species. The alignment comprised 1,824,530 amino-acid positions and was partitioned by gene. Model selection and partition merging were performed in IQ-TREE v3.0.1 [71] using ModelFinder Plus with the MFP+MERGE option. Branch support was assessed using 1,000 ultrafast bootstrap replicates. To assess phylogenetic structure in chromosome-scale synteny conservation, a species-level synteny score was calculated as the mean chromosome mapping proportion across all pairwise comparisons involving each species as reference. These scores were mapped onto the phylogeny, and phylogenetic signal was quantified using Blomberg’s K. No autosome–sex chromosome synteny links were detected among species with reliably identified sex chromosomes under the applied homology classification and filtering criteria, and these comparisons were therefore not analyzed further. Differences between autosome–autosome and sex chromosome–sex chromosome comparisons were tested using a linear mixed-effects model, with chromosome-level synteny mapping proportion as the response variable and chromosome comparison class as the fixed predictor. Reference and target species were included as random intercepts to account for non-independence among chromosome comparisons involving the same species. Statistical significance was further evaluated using permutation tests. Chromosome homology-group stability was additionally assessed by quantifying the frequency with which chromosomes retained consistent homology assignments across pairwise species comparisons.

Next, structural chromosome conservation was assessed by integrating chromosome length data for all species. Evolutionary size divergence between homologous chromosomes was quantified as the absolute log-transformed difference in chromosome length between reference and target chromosomes. Differences in chromosome size divergence between autosomes and sex chromosomes were tested using a linear mixed-effects model, with chromosome size divergence as the response variable and chromosome comparison class (autosome–autosome vs. sex chromosome–sex chromosome) as the fixed effect. Reference species and target species were included as crossed random intercepts to account for non-independence among chromosome comparisons, as the same species contributed to multiple pairwise comparisons. Statistical significance was additionally evaluated using permutation tests to determine whether observed differences in structural divergence exceeded expectations based on the chromosome comparison structure.

Ancestral evolution of sex chromosome number and total sex chromosome sequence length was then reconstructed using the species-level phylogeny. Sex chromosome number was defined as the total number of chromosomes assigned to sex-linked homologous groups based on the homology-based classification described above. The discrete trait was reconstructed using corHMM, comparing equal-rates (ER), all-rates-different (ARD), and ordered (STEP) transition models. The ER model had the lowest AIC and was therefore used for ancestral-state reconstruction. For each species, total sex chromosome sequence length was calculated as the sum of the lengths of all chromosomes assigned to sex-linked homologous groups. This continuous trait was log-transformed as log(total sex chromosome sequence length + 1), and ancestral values were estimated using maximum-likelihood reconstruction under a Brownian-motion model implemented in phytools::fastAnc. The resulting continuous trait was mapped onto the phylogeny using phytools::contMap. These reconstructions were used to characterize the evolutionary history of sex chromosome number and total sex chromosome sequence length across the sampled spider lineages.

Finally, to test whether sex chromosome size scales with overall genome size or genome-wide repeat content, species-level linear regression analyses were performed using total sex chromosome length, genome size, and repeat fraction estimates derived from chromosome-scale genome assemblies. Separate linear models were fitted to evaluate relationships between (i) sex chromosome size and genome size, (ii) sex chromosome size and repeat fraction, and (iii) genome size and repeat fraction.

## Results

### Genome survey and draft assembly

The *Nephilingis cruentata* genome was initially sequenced with MGI short reads (109 Gbp) to conduct a genome survey. After quality filtering, 101.3 Gbp (Q20: 99.43%) of short reads with a GC content of 32.02% were retained for calculating k-mer based statistics. KMC-based k-mer analyses calculated a genome size of 1,900,753,108 bp with a heterozygosity of 1.30% (**Table S1**).

Based on these genome survey statistics, PacBio HiFi long reads (73.4 Gbp) were sequenced (**Table S1; Figure S1**). After quality filtering, HiFi reads (mean length 17.7 kbp, N50 17.7 kbp) were used for a long-read only assembly with hifiasm, followed by redundancy removal. The preliminary assembly had a span of 1.72 Gbp, with a contig N50 of 72.63 Mbp and the longest contig reaching 145.6 Mbp. The assembly comprised 233 contigs ≥1 kbp (**Table S6**).

Genome completeness was evaluated using BUSCO with the arachnida_odb10 dataset, identifying 97.96% complete BUSCOs (93.29% single-copy, 4.67% duplicated; **Table S3**). K-mer analysis showed a QV of 58.43 with an estimated consensus error rate of 0.0000014, and completeness values of 81–87% depending on input data type. Mapping accuracy was high: 99.83% of NGS reads and 99.99% of HiFi reads aligned back to the assembly. Coverage depth was ∼58× for NGS, and ∼42× for HiFi reads, with an average GC content of 31% (**Figure S2**). After removal of minor contaminant sequences, the assembly size remained 1.72 Gb with unchanged contiguity statistics.

### Hi-C chromosome-scale assembly

To achieve chromosome-scale resolution, 244 Gbp of Hi-C data were generated and quality-filtered, yielding 1.60 billion clean reads (clean rate 98.2%, Q30 89.7%). After alignment and filtering, 185,996 valid Hi-C interaction pairs were retained (3.3% of clean reads; 67.8% of uniquely mapped reads).

Using HapHiC, which integrates 3D-DNA and ALLHiC-based ordering and orientation algorithms, 1.71 Gbp (99.45%) of the sequence data was anchored to 13 pseudochromosomes, corresponding to the expected haploid karyotype (**Table 1**). The resulting assembly reached a scaffold N50 of 131.6 Mbp, with the longest scaffold (Chr01) spanning 145.6 Mbp. The final assembly comprised 13 pseudochromosomes, with a total of 194 scaffolds and 36 gaps (**Table 1; Table S7**).

**Table 1.**
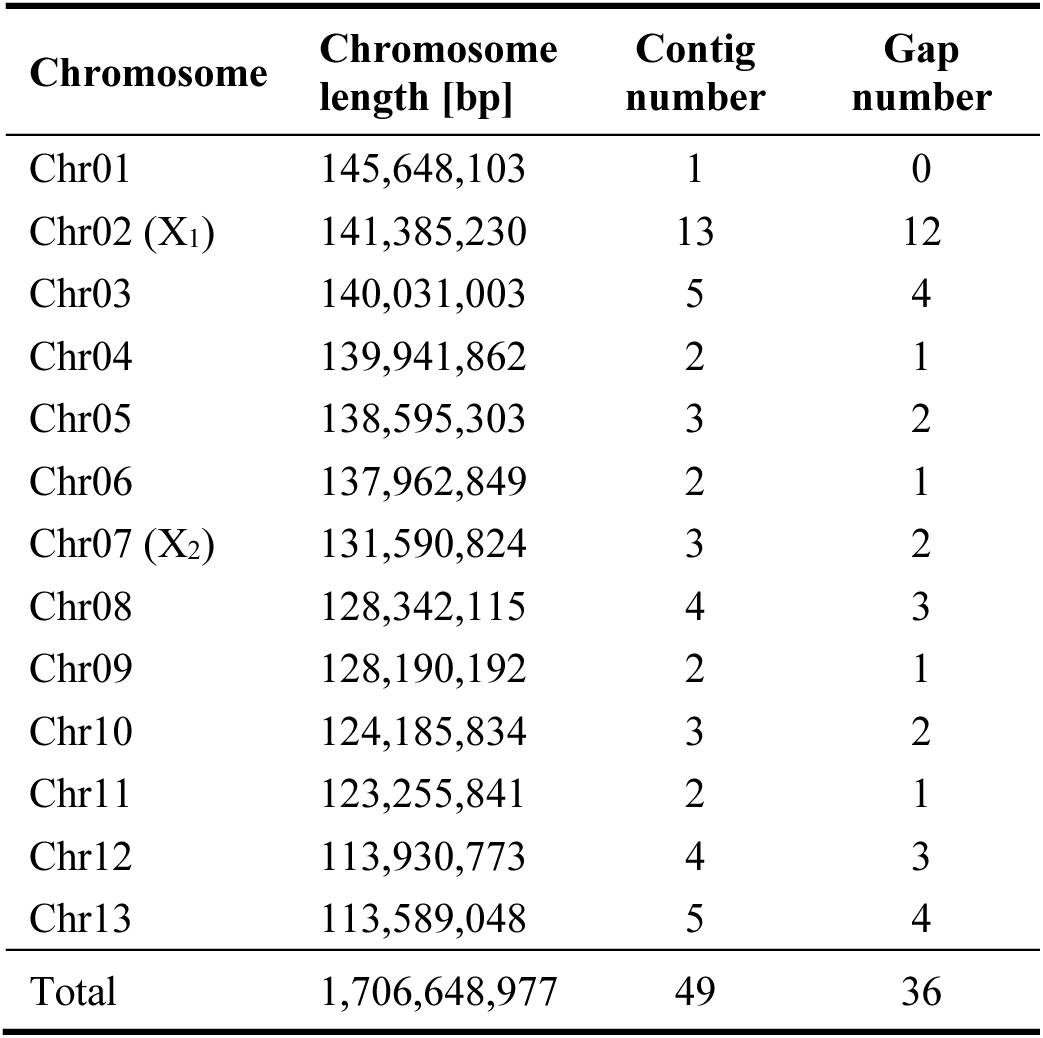
Chromosome-scale pseudochromosomes generated through Hi-C scaffolding. Two sex chromosomes (X_1_ and X_2_) were identified based on reduced sequencing coverage in males relative to females. The final assembly comprised 13 pseudochromosomes with a combined length of 1.71 Gbp. For each pseudochromosome, chromosome length, contig number, and gap number are shown.

Interaction heatmaps showed strong intra-chromosomal contact enrichment and minimal inter-chromosomal noise, confirming accurate clustering and orientation (**Figure 1A**). Genome completeness improved further, with 98.8% complete BUSCOs detected (96.2% single-copy, 2.7% duplicated) (**Figure 1B; Table S3**).

**Figure 1.**
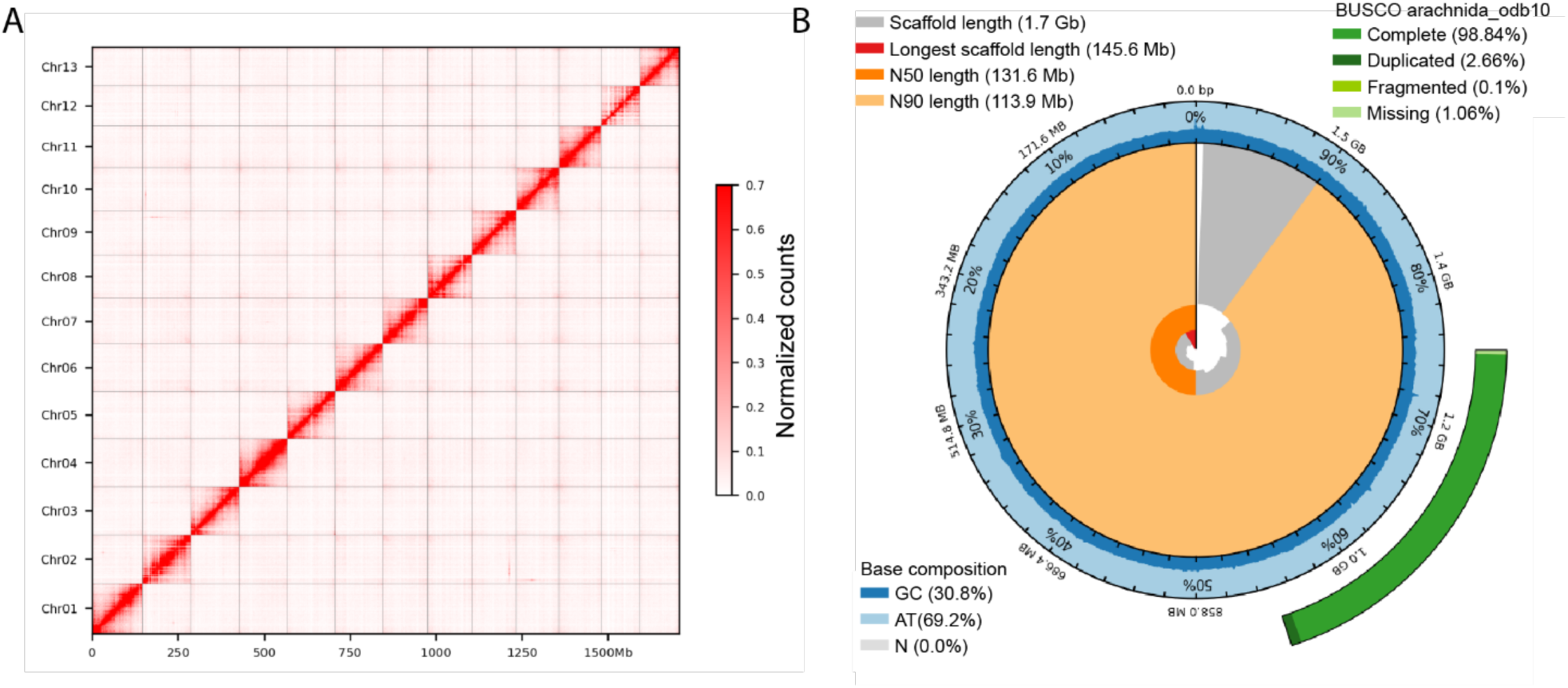
Chromosome-scale assembly of the *Nephilingis cruentata* genome. Chromosome-scale genome assembly of the *Nephilingis cruentata* supported by Hi-C scaffolding (A) and standard assembly statistics (B). The Hi-C contact map resolves 13 chromosome-scale scaffolds, consistent with the inferred karyotype, with clear compartmentalisation of interaction signals indicating strong contiguity. Assembly quality is summarised in a snail plot integrating key indicators of genome completeness, contiguity, and sequence composition. Together, these metrics show a highly complete and well-resolved genome assembly with strong support for chromosome-level reconstruction.

### Repeat and gene annotation

The final chromosome-scale assembly (1.72 Gbp; scaffold N50 = 131.6 Mbp) was annotated using combined transcriptome-based, homology-based, and de novo approaches. Repetitive elements accounted for 42.7% of the genome, dominated by transposable elements (39.8% of the genome). DNA transposons represented the largest fraction (21.1% of the genome), followed by LTRs (6.9%), LINEs (6.5%), and SINEs (2.3%). Simple sequence repeats (SSRs) were also abundant, with 245,237 loci identified (**Table S8**).

After integration of gene models with EvidenceModeler, the genome annotation included 20,021 predicted protein-coding genes, with an average gene length of 27.7 kbp, an average CDS length of 1,468 bp, and ∼7.2 exons per gene (**Table S9**). Functional annotation was assigned to 95.8% of predicted genes, indicating that the majority of predicted proteins were assigned putative functions (**Table S10**).

### Identification of candidate X chromosomes using sex-specific coverage

Normalized chromosome-level coverage was broadly consistent across autosomes, which clustered close to the expected baseline of 1.0 in both males and females (**Figure 2A**). In contrast, two chromosomes (Chr02 and Chr07) showed consistent deviations from this autosomal baseline. In males, these chromosomes exhibited reduced normalized coverage relative to the genome-wide expectation, while female coverages for these chromosomes remained close to autosomal baseline (compared to male values) (**Figure 2A**). This pattern is also reflected in the direct comparison between sexes, where Chr02 and Chr07 show the strongest negative deviations in male–female differences (Fig. 2B). The resulting male-to-female coverage ratios for these chromosomes were approximately 0.62–0.64, clearly separated from the distribution of autosomes, which clustered around a ratio of ∼1.0. Together, these coverage patterns identify Chr02 and Chr07 as candidate X chromosomes, while all other chromosomes show no evidence of sex-biased coverage differences.

**Figure 2.**
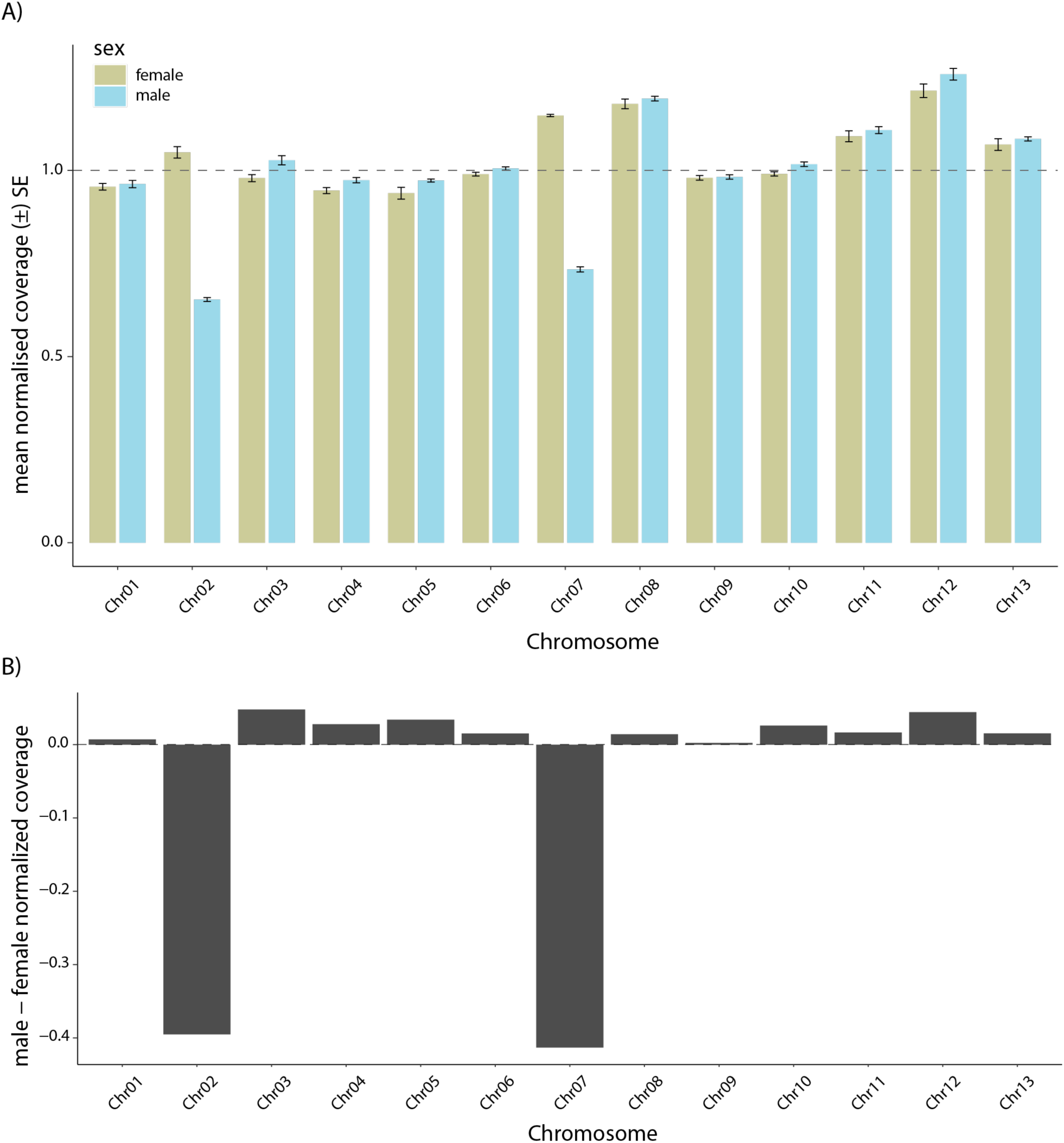
Identification of sex chromosomes (X1 and X2) in *Nephilingis cruentata*. Normalized chromosome-level coverage is broadly similar between males and females for most chromosomes, clustering around a common autosomal baseline (∼1.0). In contrast, Chr02 and Chr07 show consistent reductions in male relative coverage compared to females. Sex-specific normalized coverage per chromosome is shown (A), and corresponding male–female differences in normalized coverage highlight the strongest deviations for Chr02 and Chr07 (B).

### Stability of sex chromosome evolution in spiders

Comparative synteny analysis across chromosome-scale spider genomes revealed that chromosomes cluster into distinct homology groups, with sex chromosomes forming a consistent subset of these clusters (**Table S11**). All experimentally identified and homology-inferred sex chromosomes were assigned to a single sex-linked homology group, indicating strong conservation of sex chromosome identity across spider lineages (**Figure 3A**). Despite this conserved chromosomal identity, sex chromosomes exhibited significantly reduced synteny conservation compared with autosomes (**Figure 3B**). Linear mixed-effects modelling showed a strong effect of chromosome class on synteny weight (*F_1,1296_* = 214.97, *P* < 0.0001), with sex chromosome comparisons displaying substantially lower mean synteny conservation than autosomal comparisons (0.583 vs. 0.733). Nevertheless, sex chromosome comparisons remained entirely restricted to the same inferred homology groups (stability = 1.0), whereas autosomal comparisons showed lower overall homology stability (0.869), indicating that sex chromosomes retain conserved chromosomal identity despite elevated sequence and structural divergence. Phylogenetic analyses revealed only weak evidence for phylogenetic structuring of synteny conservation across species (Blomberg’s *K* = 1.12, *P* = 0.071), suggesting that phylogenetic relatedness alone does not explain most variation in synteny conservation across the sampled taxa. Chromosome size evolution analyses showed elevated divergence of sex chromosomes relative to autosomes (**Figure 3B**). Homologous sex chromosome pairs showed significantly greater log-transformed chromosome length divergence than homologous autosomal pairs (0.827 vs. 0.613; linear mixed-effects model: *F_1,1129_* = 54.91, *P <* 0.0001). This pattern remained significant when analyses were restricted to species possessing exactly two inferred sex chromosomes (**Figure 3B**), indicating that the result is not driven by differences in sex chromosome copy number among species (*F_1,816_* = 30.99, *P <* 0.0001).

**Figure 3.**
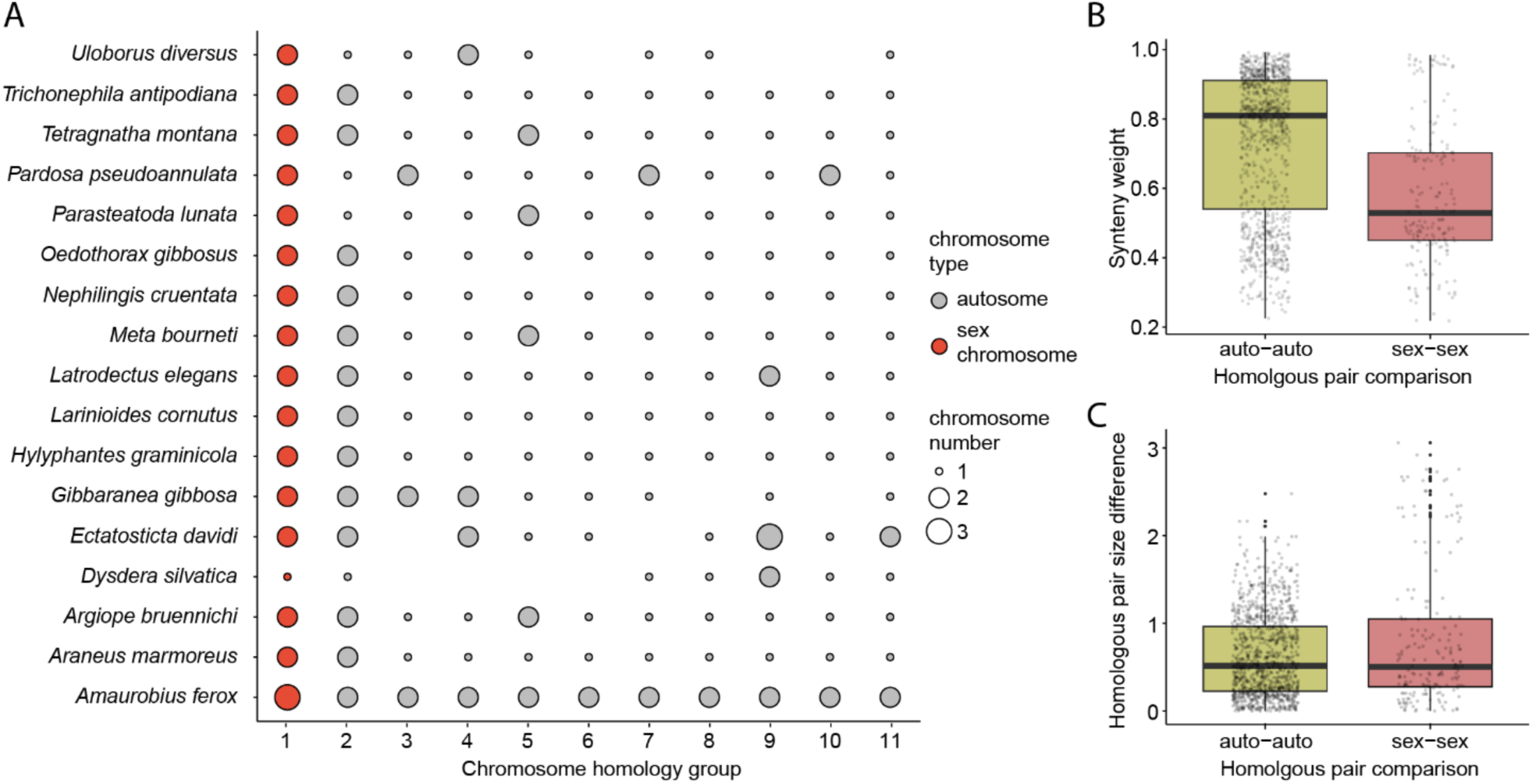
Conserved sex chromosome homology despite elevated structural divergence across spider species. (A) Chromosome homology groups across the sampled species, showing that chromosomes assigned as sex chromosomes consistently cluster within a single homology group, despite variation in the number of sex chromosomes among species. Point size indicates the number of chromosomes assigned to each homology group, and colours distinguish sex chromosomes from autosomes. (B) Comparison of synteny mapping weights between autosome– autosome and sex chromosome–sex chromosome homologous pairs, showing generally higher conservation in autosomal pairs. (C) Distribution of absolute chromosome length differences among homologous pairs, indicating greater structural divergence between sex chromosome pairs compared to autosomal pairs.

Ancestral state reconstruction of sex chromosome copy number supported variation in inferred sex chromosome counts across the phylogeny (**Figure 4A**). Comparative model fitting favoured an equal-rates model of chromosome number evolution over asymmetric or constrained stepwise alternatives (AIC*_ER_* = 19.72; AIC*_STEP_* = 20.98; AIC*_ARD_* = 27.84), suggesting relatively simple transition dynamics across the sampled taxa. Continuous trait reconstruction under a Brownian-motion model further revealed substantial interspecific variation in total sex chromosome length across the phylogeny (**Figure 4B**).

**Figure 4.**
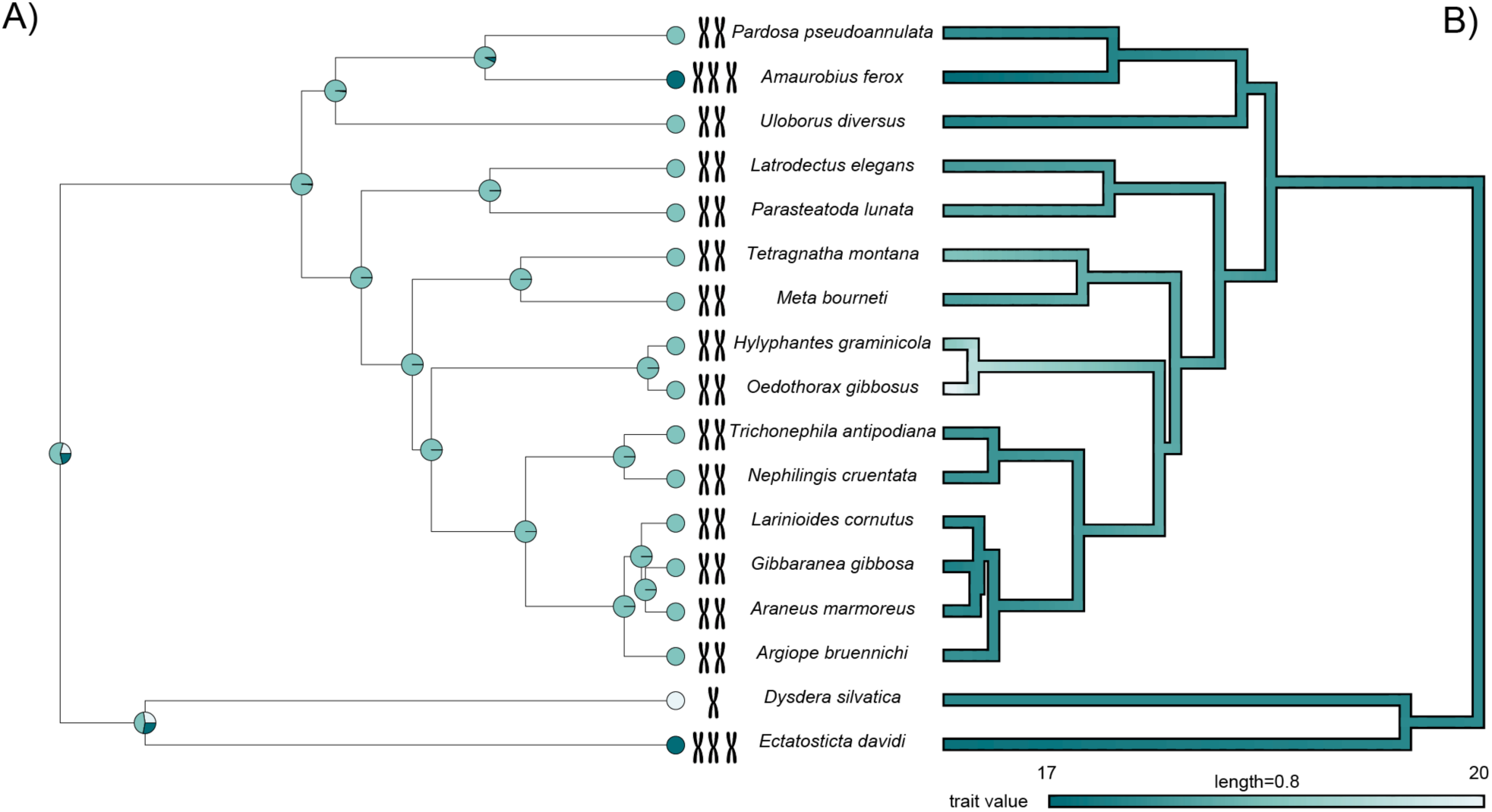
Ancestral trait reconstruction of sex chromosome number and size in spiders. Ancestral states of sex chromosome number were reconstructed from sex-linked chromosome assignments based on available annotations and synteny-informed homology (A). Among the sampled taxa, sex chromosome number is relatively conserved, with most species exhibiting an X_1_X_2_ system. Total sex chromosome sequence length, calculated as the summed length of inferred sex chromosomes, is shown on a logarithmic scale (B). Sex chromosome sequence length varies substantially among lineages, with species possessing three sex chromosomes showing particularly large total sex chromosome sequence length

Across species, total sex chromosome size was strongly positively associated with overall genome size (*F_1,15_* = 50.81, *R²* = 0.77, P < 0.0001), indicating that species with larger genomes generally also possess more and/or larger sex chromosomes. A significant relationship was found between sex chromosome size and genome-wide repeat fraction (*F_1,15_* = 10.001, *R²* = 0.40, *P* = 0.006). Similarly, total genome size was correlated with genome-wide repeat fraction (*F_1,15_* = 8.167, *R²* = 0.35, *P* = 0.012), albeit the relationship was a bit weaker. A direct comparison of the two dependent correlations did not provide evidence that these associations differed significantly (Steiger’s test: *z* = 0.51, *p* = 0.607). These results suggest that larger spider genomes are in part driven by increased repetitive sequence content.

## Discussion

Spiders provide a strong system for studying the genomic basis of phenotypic evolution, with many lineages showing substantial variation in silk and venom biology, reproductive biology, life history, sexual size dimorphism, and sex chromosome systems [1–3,14]. However, the genomic architecture underlying many of these traits remains poorly resolved, particularly at the chromosome scale [e.g. 5,9,72]. The African hermit spider *Nephilingis cruentata* is a particularly informative case: females are approximately 75 times heavier than males, development differs strongly between the sexes, and reproduction involves mating plugs, pedipalp breakage, and sexual cannibalism [25–27,29]. Cytogenetic work has established an X₁X₂ system in this species [28], but the corresponding chromosomes had not yet been identified in a chromosome-level genome assembly. Here, we provide this genomic framework, with 99.5% of bases assigned to 13 pseudochromosomes, a scaffold N50 of 131.6 Mbp, and a BUSCO completeness score of 98.8%. We identify Chr02 and Chr07 as candidate X chromosomes and show that, across the analysed chromosome-scale spider genomes, sex chromosomes retain broad homologous identity while exhibiting lower synteny conservation and greater chromosome-length divergence than autosomes.

Sex-specific whole-genome resequencing allowed us to assign the known X₁X₂ system of *N. cruentata* to specific pseudochromosomes in the assembly. By sequencing five males and five females at approximately 10× coverage and comparing normalized chromosome-level read depth, we found that most chromosomes clustered around the expected autosomal baseline in both sexes, whereas Chr02 and Chr07 showed a consistently reduced relative coverage in males. This pattern is expected for X-linked chromosomes under male hemizygosity and identifies Chr02 and Chr07 as candidate genomic representatives of X₁ and X₂. Comparable genomic and cytogenetic approaches have been used to identify sex chromosomes in other chromosome-level spider genomes, including *Argiope bruennichi, Uloborus diversus,* and *Latrodectus hesperus* [6,23,73]. Thus, the *N. cruentata* result adds to the established approaches for detecting sex chromosomes in spiders, while providing the first chromosome-level assignment of the inferred X₁X₂ system in Nephilidae. However, identifying candidate X chromosomes is not the same as identifying the molecular mechanism of sex determination. Resolving recombination patterns, candidate sex-determining loci, and possible sex-linked regulatory mechanisms will require higher-coverage resequencing, cytogenetic validation, and sex-specific expression analyses.

The main comparative result across the analysed spider lineages is that chromosomes assigned to sex-chromosome homology groups retain stable chromosomal identity across the sampled taxa. This finding is relevant to a long-standing unresolved question in spider sex chromosome evolution: the origin of the ancestral X₁X₂ system. The X₁X₂ system has been proposed to have evolved from an ancestral X₀ system through fission of a single X chromosome, implying a common evolutionary origin of the two X chromosomes [13,74]. Alternatively, X₁ and X₂ have been proposed to represent distinct, non-homologous sex chromosomes [74,75]. Our finding that sex chromosomes consistently occupy the same broad homology groups across the sampled lineages is consistent with the hypothesis of conserved chromosomal identity and therefore provides comparative genomic support for a shared evolutionary origin of spider X chromosomes (**Figure 3A**). Importantly, however, this conserved chromosomal identity is accompanied by substantial structural divergence: sex chromosomes show lower synteny conservation and greater chromosome-length divergence than autosomes (**Figure 3B, C**). Although cytogenetic studies have emphasized the lability of spider sex chromosome systems, including variation in chromosome number and the occurrence of fissions, fusions, nondisjunctions, and sex chromosome–autosome rearrangements [13,14], our comparative genomic analyses indicate that such variation does not necessarily require repeated replacement of the underlying sex-linked chromosome material. Instead, the consistent recovery of X-chromosome homologues within the same homology groups across sampled spider lineages suggests that sex chromosome identity can be evolutionarily conserved while chromosome structure undergoes lineage-specific remodeling. Similar patterns, in which conserved sex chromosome ancestry coexists with substantial structural divergence, have also been reported in other taxa, indicating that chromosome identity and chromosome architecture can evolve at different rates [76,77]. However, sex chromosome evolution is highly diverse across animals, with some groups showing long-term conservation of ancestral sex chromosomes, whereas others exhibit repeated sex chromosome turnover and recruitment of new genomic regions into sex-determining roles [19,78]. Our results add comparative genomic evidence for a shared evolutionary origin of spider sex chromosomes and demonstrate that, within this conserved chromosomal framework, spider sex chromosomes have undergone accelerated structural evolution relative to autosomes. Thus, spiders provide another example of the diverse evolutionary trajectories through which sex chromosome systems can arise and diversify across animal lineages.

Although weak phylogenetic signal may partly reflect limited and uneven taxon sampling, variation in synteny conservation was not strongly predicted by phylogenetic relatedness across the sampled taxa. Instead, the stronger structural divergence of X-chromosome homology groups relative to autosomes likely reflects lineage-specific processes affecting chromosome organization. One plausible route is recombination suppression: once recombination is reduced or halted on sex chromosomes, constraints maintaining gene order may be weakened, allowing inversions and other rearrangements to persist more readily; alternatively, inversions themselves may contribute to recombination suppression [20]. Fissions, fusions, segmental gains and losses, and repeat-mediated changes may further contribute to differences in synteny conservation and chromosome length. These mechanisms are consistent with broader models of sex chromosome evolution, in which sex-specific transmission, altered effective population size, recombination suppression, and sexually antagonistic selection can cause sex chromosomes to diverge from autosomes [19–21,79]. However, our analyses do not directly identify the causal drivers of this divergence. Because our results are based on chromosome-scale synteny conservation and chromosome-length divergence rather than dN/dS, substitution rates, or protein evolution, they should be interpreted as evidence for elevated structural lability of X-chromosome homology groups, not as evidence for classical faster-X sequence evolution. Testing whether spider X chromosomes also show faster molecular evolution will require comparative analyses of X-linked and autosomal coding sequences, regulatory regions, and sex-biased expression.

The assignment of Chr02 and Chr07 as the X-linked pseudochromosomes corresponding to X₁ and X₂ in *N. cruentata* places previous work on sexual size dimorphism into a chromosomal context. Quantitative-genetic analyses showed that adult body size has a sex-specific architecture, with female size mainly associated with direct genetic effects and male size more strongly associated with maternal effects [25]. Although the present assembly does not directly contribute to explaining the evolution of extreme female-biased sexual size dimorphism, it establishes the chromosomal framework critically needed to distinguish autosomal, X-linked, and structurally divergent genomic contexts for sex-specific traits. In this sense, the current study connects existing phenotypic and quantitative-genetic knowledge by providing a chromosome-level resource for studying sex-specific development, reproductive traits, and phenotypic variation at the genomic level in *N. cruentata*.

In conclusion, the *N. cruentata* reference genome provides a chromosome-scale resource for Nephilidae and a comparative framework for studying sex chromosome evolution in spiders. The results suggest that spider sex chromosomes in the sampled assemblies are not primarily labile in homologous identity, but in structure. This distinction helps reconcile cytogenetic diversity with genomic conservation: X chromosome number and size may vary, while sex-linked chromosomal material remains broadly conserved. For *N. cruentata*, the assembly provides the chromosomal framework needed to place sex-specific development, female-biased size, and reproductive traits in relation to X-linked, autosomal, and structurally divergent genomic regions.

## Acknowledgments

We thank Behare Rexhepi, Nik Lupše, and Lucija Kunavar for their help in the molecular laboratory, and C. Haddad, M. Gregorič, K. Čandek, S. G. Quiñones-Lebrón, T. Lokovšek, and M. Kuntner for collecting founder individuals in South Africa that were used to establish the laboratory population (permits OP 552/2015 and OP 3031/2020).

