## Supplemental Figures for "High-quality reference genome of the African hermit spider, *Nephilingis cruentata,* and sex chromosome evolution in spiders"

**Additional files**

**Tables S1–S11** are provided in a separate Excel file.

**Figure S1. PacBio HiFi read length distribution.** PacBio HiFi read lengths show a sharp peak around ~12 kb, followed by a gradual decline extending to ~45 kb. The distribution is right-skewed with a long tail of longer reads.

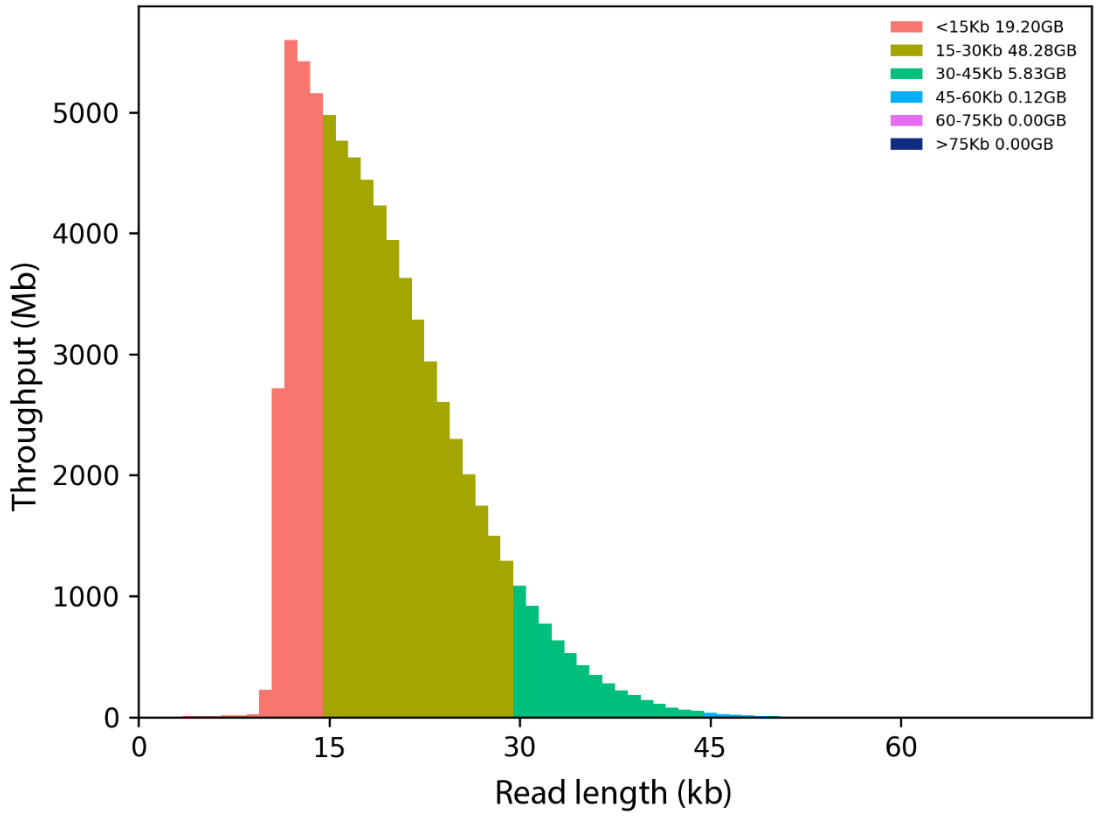

**Figure S2. PacBio HiFi GC content and sequencing depth distribution.** A density plot was generated from third-generation read alignments to the assembled genome, summarizing GC content and sequencing depth. GC content is distributed between 25–35%, while sequencing depth is concentrated around 30–50×.

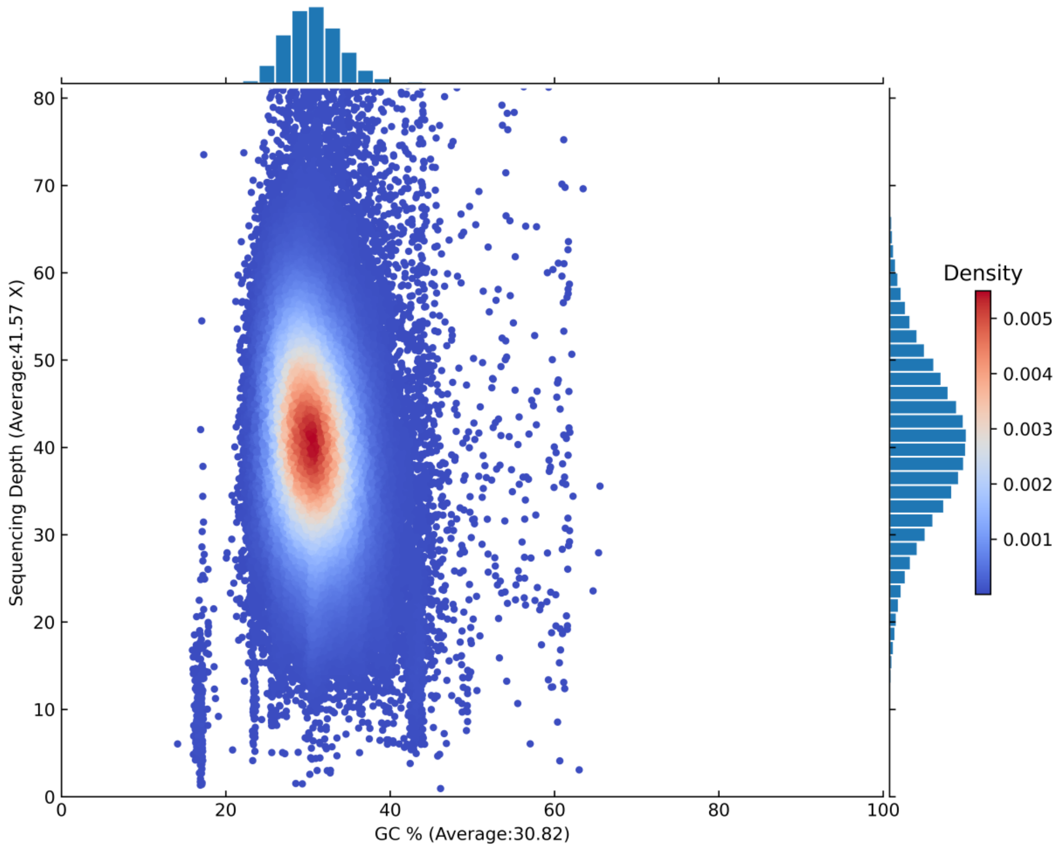

**Figure S3. Relationship between genome size, X chromosome size and repeat content.** Sex chromosome size positively covaries with genome size (A), meaning that larger genomes also tend to have larger sex chromosomes. Sex chromosome size (B) and genome size (C) further correlates with repeat content of the genome. This shows that larger sex chromosomes and larger genomes in general tend to contain more repetitive sequences, accounting for around 35% of the variation overall.

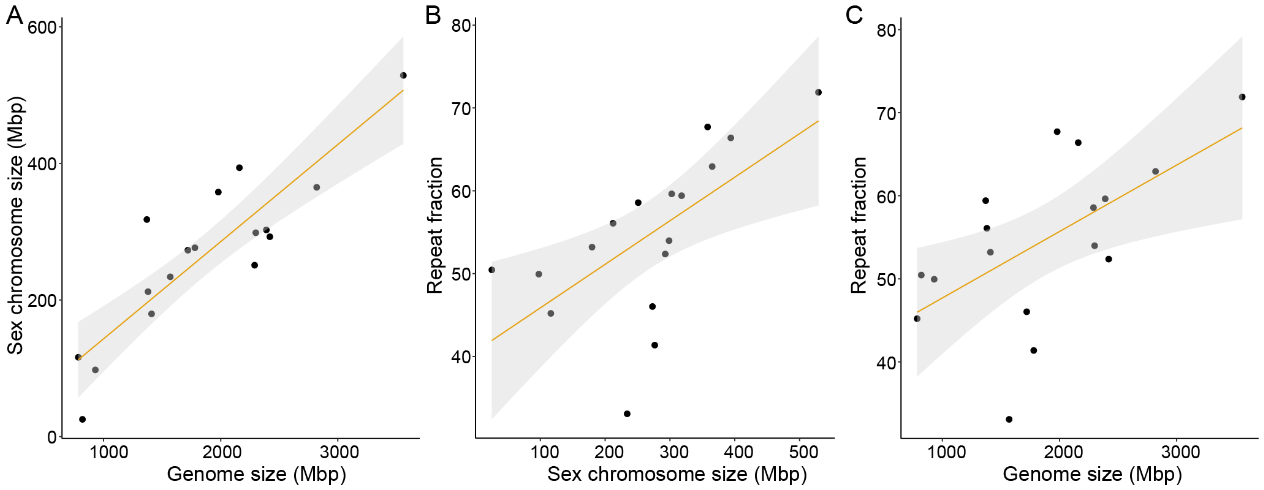
